# Plant-soil feedback contributes to predicting plant invasiveness of 68 alien plant species differing in invasive status

**DOI:** 10.1101/568048

**Authors:** Anna Aldorfová, Pavlína Knobová, Zuzana Münzbergová

**Affiliations:** Department of Botany, Faculty of Science, Charles University, Benátská 2, 128 01 Prague 2, Czech Republic; Institute of Botany, Czech Academy of Sciences, Zámek 1, 252 43 Průhonice, Czech Republic

**Keywords:** alien / exotic / non-native species, enemy release hypothesis, intraspecific (conspecific) plant-soil feedback, invasive ecology, neophytes, plant–soil (below-ground) interactions, phylogenetic relatedness, residence time, specific leaf area.

## Abstract

1. Understanding what species characteristics allow some alien plants to become invasive while others fail to is critical to our understanding of community assembly processes. While many characteristics have been shown to predict plant invasiveness, the importance of plant-soil feedbacks (PSFs) in invasions has been difficult to assess since individual studies include only a few species and use disparate methodology.

2. We studied PSFs of 68 invasive and non-invasive alien species in a single two-phase garden experiment, and compared the relative importance of PSF, residence time, phylogenetic novelty and plant traits for plant invasiveness. Additionally, we tested for relationships between PSF, residence time and phylogenetic novelty.

3. PSF for seedling establishment belonged to five best predictors of plant invasiveness, along with specific leaf area, height, seedling growth rate, and residence time. Invasive species had more positive PSF for seedling establishment, but not for biomass, than non-invasive species. Phylogenetically novel species experienced less negative PSF than species with native congeners, suggesting they benefit more from enemy release. PSF of non-invasive species, contrary to that of invasive species, was becoming more negative with increasing residence time.

4. ***Synthesis***. We demonstrated that PSF plays a role in predicting invasiveness that is comparable with other species characteristics that are more commonly studied. PSF should thus receive more attention in studies predicting community structure and in programs assessing the likely invasions of aliens.

## Introduction

Understanding the causes of biological invasions is a priority of ecological research in the last decades (Sol et al., 2012). Invasive plants displace native species, change vegetation structure, reduce native biodiversity (Hejda et al., 2009, Powell et al., 2013), undermine functioning of the whole ecosystems (Richardson and Pysek, 2012) and cause significant economic losses (Zavaleta, 2000). It is therefore in the interest of our society to determine the causes of species invasiveness and to prevent the emergence of new invasive species.

Many previous studies have examined species characteristics that promote plant invasiveness and differentiate invasive species from their non-invasive relatives. Common characteristics that have been considered, and showed importance in some systems, are plant traits (Rejmanek and Richardson, 1996, van Kleunen et al., 2010), phylogenetic novelty (Strauss et al., 2006), residence time (Pysek and Jarosik, 2005) or plant-soil feedback (Callaway et al., 2004, Kulmatiski et al., 2008, Klironomos, 2002). However, studies only rarely compare the predictive power of multiple species characteristics (Lowry et al., 2013), which allows their relative importance to be ranked, and none of the comparisons have so far included information on plant-soil feedback.

Intraspecific plant-soil feedback (PSF) has been suggested to play an important role in plant invasions. Invasive species generate more positive or less negative intraspecific PSF than native species (e.g., van Grunsven et al., 2007, Kulmatiski et al., 2008, Klironomos, 2002, Meisner et al., 2014). Invasive species also often experience less negative intraspecific PSF in their introduced range compared to the native range due to enemy release (e.g., Reinhart and Callaway, 2004, Callaway et al., 2004, Maron et al., 2014, Yang et al., 2013, Gundale et al., 2014). The extent to which PSF may predict plant invasiveness, i.e. determine the invasion success of alien plants, however, remains unclear. Alien species differ in the degree of enemy release as well as in the way they interact with individual components of soil biota. Some alien species are therefore likely to develop less negative or more positive PSF than other alien species, and thus are more likely to become invasive. A synthesis on this topic is lacking due to limited number of alien non-invasive species that have been studied and due to the disparate methodology across studies. A meta-analysis of Kulmatiski et al. (2008) suggested that invasive species create less negative PSFs than non-invasive alien species. Their analysis was, however, based on only a few non-invasive aliens studied in independent experiments and all occurring in grassland ecosystems. The results thus need to be confirmed in a single study with more species, consistent methodology and with species from other ecosystems.

The strength of intraspecific PSFs is also expected to be related to other species characteristics than invasive status, such as residence time and phylogenetic novelty of the focal alien. PSFs might become more negative with increasing residence time as local pathogens colonize or adapt to the focal alien (Dostal et al., 2013, Flory and Clay, 2013). The literature has shown mixed support for this idea, with one study showing increasingly more negative intraspecific PSF with residence time of aliens (Diez et al., 2010) and two others showing no effect (Speek et al., 2015, McGinn et al., 2018). One of the reasons for the disparity across studies might be that the studies did not consider the invasive success of the aliens. The expected negative pattern might be particularly pronounced for non-invasive aliens that get regulated by the accumulating pathogens while the invasive species might be less prone to pathogen accumulation and thus be able to attain dominance (Kulmatiski et al., 2008).

PSF should be more negative for aliens that are closely related to plants in the native flora because related plant species are more likely to share pathogens (Bufford et al., 2016, Vacher et al., 2010, Parker et al., 2015, Gilbert and Parker, 2016). While it has been shown that PSFs are phylogenetically conservative, i.e. that close relatives exhibit more similar feedback responses than phylogenetically distant species (Anacker et al., 2014), no studies examined the effect of phylogenetic novelty of aliens on their intraspecific PSFs. However, research on interspecific PSFs (experiments that compare plant fitness in soil conditioned by the focal plant and soil conditioned by another plant species) demonstrated that plants experience more negative interspecific PSF when grown in soil conditioned by a congener than in soil conditioned by a less related species (Callaway et al., 2013, Dostal and Paleckova, 2011, Kempel et al., 2018, but see Kutakova et al., 2018, Munzbergova and Surinova, 2015, Mehrabi and Tuck, 2015).

In this study, we quantified intraspecific PSF of 68 alien species of the Czech Republic that vary in their invasive status, and abundance and dominance in the field. We evaluated the direct and interactive effects of invasive status, residence time and phylogenetic novelty on PSFs. Moreover, we compiled data on a broad range of plant traits (seven whole plant traits, five regenerative traits and one leaf trait) from trait databases, previous studies using the same set of species (Kubesova et al., 2010, Moravcova et al., 2010) and our own data collection, and ranked the plant traits, PSF, minimum residence time and phylogenetic novelty according to their importance for predicting invasiveness of our focal species. The hypotheses we tested were as follows: i) Invasive species have less negative or more positive intraspecific PSF than non-invasive alien species; ii) Phylogenetically novel species have less negative or more positive PSF than species with a close native relative; iii) PSF is becoming more negative with increasing residence time and the effect is more pronounced for non-invasive species than for invasive species; iv) PSF is as important for predicting plant invasiveness as some other commonly used traits.

## Materials and methods

### Studied species

In this study, we used 68 neophytes (alien species introduced after 1500 A.D. sensu Pysek et al. (2004)) occurring in the Czech Republic (Table S1). The species were selected based on a list of 93 species used in previous studies of Moravcova et al. (2010) and Kubesova et al. (2010) who compared reproductive characteristics and genome size of alien invasive and non-invasive species, respectively. The original set of species was reduced because i) some species from the list were reclassified to archeophytes by Pysek et al. (2012) and ii) we did not manage to collect seeds or cultivate some of the species. Choosing species from that list allowed us to determine the relative importance of PSF and many other traits studied previously for plant invasiveness.

Invasion status (casual, naturalized non-invasive or invasive, sensu Pysek et al. (2004)) of the studied species in the Czech Republic was taken from Pysek et al. (2012). Invasive species in our study are understood as species that form self-replacing populations over many life cycles, produce reproductive offspring, often in very large numbers, at considerable distances from the parent and/or site of introduction, and have the potential to spread over long distances (Pysek et al., 2004). Of the 68 species, 27 neophytes are considered invasive in the Czech Republic and 41 non-invasive. The vast majority of the non-invasive species are classified as naturalized non-invasive (i.e. forming self-sustaining populations, not depending on human intervention, but not spreading (Pysek et al., 2004)) and only three (*Ambrosia trifida, Rudbeckia hirta* and *Sedum rupestre*) as casual (i.e. not forming self-sustaining populations, depending on repeated introductions of propagules (Pysek et al., 2004)). We therefore did not distinguish between non-invasive naturalized and casual species in the analyses and merged them all into one category of alien non-invasive species. For each species, we also recorded their frequency (i.e. number of occupied grid cells) and maximum cover of the species in the field (taken from the Pladias database).

The 68 species analyzed are a highly representative sample, making up 17 % of the total number of 408 naturalized non-invasive neophytes and 44 % of the total number of 61 invasive neophytes of the Czech Republic (Pysek et al., 2012) of the Czech Republic. The species studied belong to 52 genera and 27 families according to the Angiosperm Phylogeny Group classification (Stevens, 2001 onwards) with Asteraceae most represented. Annuals, monocarpic perennials and polycarpic perennials are all well represented among the species, with no difference between invasive and non-invasive species (Table S1). Invasive species do not differ from the non-invasive species in terms of minimum residence time in the Czech Republic (Table S1). The invasive species have significantly higher maximum cover in the field than the non-invasive species and occupy more grid cells (Table S1). Phylogenetic novelty in terms of presence of a native congener in the Czech Republic does not differ between invasive and non-invasive species (Table S1).

### Seed collection

Seeds of all species were collected in 2014-2016 in the field from multiple localities (range 1-5, mean 2.5; Table S2) in the Czech Republic, minimum 5 km apart from each other, to account for possible differences between populations. For some species, we used seeds provided by a local commercial supplier (Planta Naturalis Ltd., Markvartice, Czech Republic) as one population. The seeds from each population were used separately in the experiments. In each population, we collected mixture of mature seeds from at least 10 individuals. Seeds were stored in paper bags under room temperature until used. The seeds were always used in the experiment in the year after their collection. The seeds of some species were cold-wet stratified for two months prior sowing (information provided by L. Moravcová) and all seeds were surface sterilized with 10% H_2_O_2_ to reduce the chance of contamination via seed surface fungi prior sowing.

### Experimental design

Following commonly used methodology (Bever et al., 1997, Kulmatiski et al., 2008), the plants were grown in a two-phase experiment. In the first (conditioning) phase, conditioned soil was prepared. In the second (feedback) phase, intraspecific PSF was studied. The experiment was carried out in the experimental garden of Institute of Botany, Czech Academy of Sciences (49°59′38.972′′N, 14°33′57.637′′E), 320 m above sea level, temperate climate zone, where the mean annual temperature is 8.6°C and the annual precipitation is 610 mm. In addition to obtaining natural rainfall, the plants were daily watered with tap water. Due to high number of studied species, the experiment was divided into three subsequent years, 2015-2017. Each year, we grew approximately the same proportion of invasive and non-invasive species.

### Conditioning phase

The aim of the conditioning phase was to prepare the soil, conditioned by the species, for the upcoming feedback phase. To set up the conditioning phase, we used a local common garden compost mixed with sand in 1:1 ratio (see Table S3 for the chemical characteristics of the soil). The garden is situated in the region where the studied species occur and the local soil thus contains the soil biota commonly encountered by the species. We prepared a homogenous heap of substrate at the beginning of the first year of the project and stored it in a dry place. To minimize possible soil heterogeneity due to different processes in the different parts of the stored soil, the soil was thoroughly mixed every time before use.

For each species and each population, we used 20 pots (10 × 10 × 10 cm) in the conditioning phase, half of which was sown with seeds of the species and the other half of the pots served as controls. Each pot with conditioned soil was randomly assigned its control pot. The pairs of pots were always kept in close proximity to each other throughout the experiment so that they were exposed to the same conditions.

Each of the ten pots per species and population was sown with ten seeds of the species. After the seeds germinated and seedlings established, we counted the seedlings and removed all but the largest one from the pot. Seedlings emerging afterwards were counted and removed. Both pots with and without plants were kept under the same conditions and regularly watered. The soil was conditioned for 12 weeks, similarly to a range of previous studies (e.g., van Grunsven et al., 2007, Meijer et al., 2011, van de Voorde et al., 2011, van Grunsven et al., 2010, Chiuffo et al., 2015, Florianova and Munzbergova, 2018). After the 12 weeks, the plants were harvested, and roots were carefully removed from the soil.

### Feedback phase

Ten seeds of a species were sown into each previously conditioned pot as well as to the control pots. For each species and each population, we thus had 10 pots with conditioned and 10 pots with control soil with sown seeds. We did not mix the soil from all the pots conditioned by the same species and population between the conditioning and feedback phase to avoid pseudoreplication (Brinkman et al., 2010).

After the seeds germinated and seedlings established, we counted the seedlings and removed all but the largest one from the pot. Seedlings emerging afterwards were counted and removed. Twelve weeks after seed germination, the plants were harvested, divided into aboveground and belowground biomass, dried to a constant weight and weighted. All plants of the same species and population were harvested from all the pots simultaneously.

### Other species characteristics affecting plant invasiveness

We compiled data on multiple species characteristics that are often related to plant invasiveness for our species from previously published papers and databases. These included minimum residence time (MRT), phylogenetic novelty, and plant traits (Fig. 4). MRT as a number of years elapsed since the first record of occurrence in the Czech Republic was taken from Pysek et al. (2012). As a measure of phylogenetic novelty, we recorded whether or not there is any species of the same genus for each focal species that is native to the Czech Republic using Danihelka et al. (2012). Plant traits considered in this study were germination, seedling growth rate, seedling establishment, propagule weight, propagule length-width ratio, buoyancy and anemochory measured as terminal velocity, taken from Moravcova et al. (2010), ploidy level and genome size, taken from Kubesova et al. (2010), and life history and releasing height, taken from the LEDA traitbase (Kleyer et al., 2008). Additionally, we measured specific leaf area (SLA) for all the species. To do this, we collected 10 leaves without leaf stalks from at least three different individuals per species prior to the harvest of the conditioning phase, dried them, weighted them and estimated the leaf area using the ImageJ program. SLA was then calculated as area/mass and an average value from the 10 measurements was used in further analyses.

**Fig. 4:**
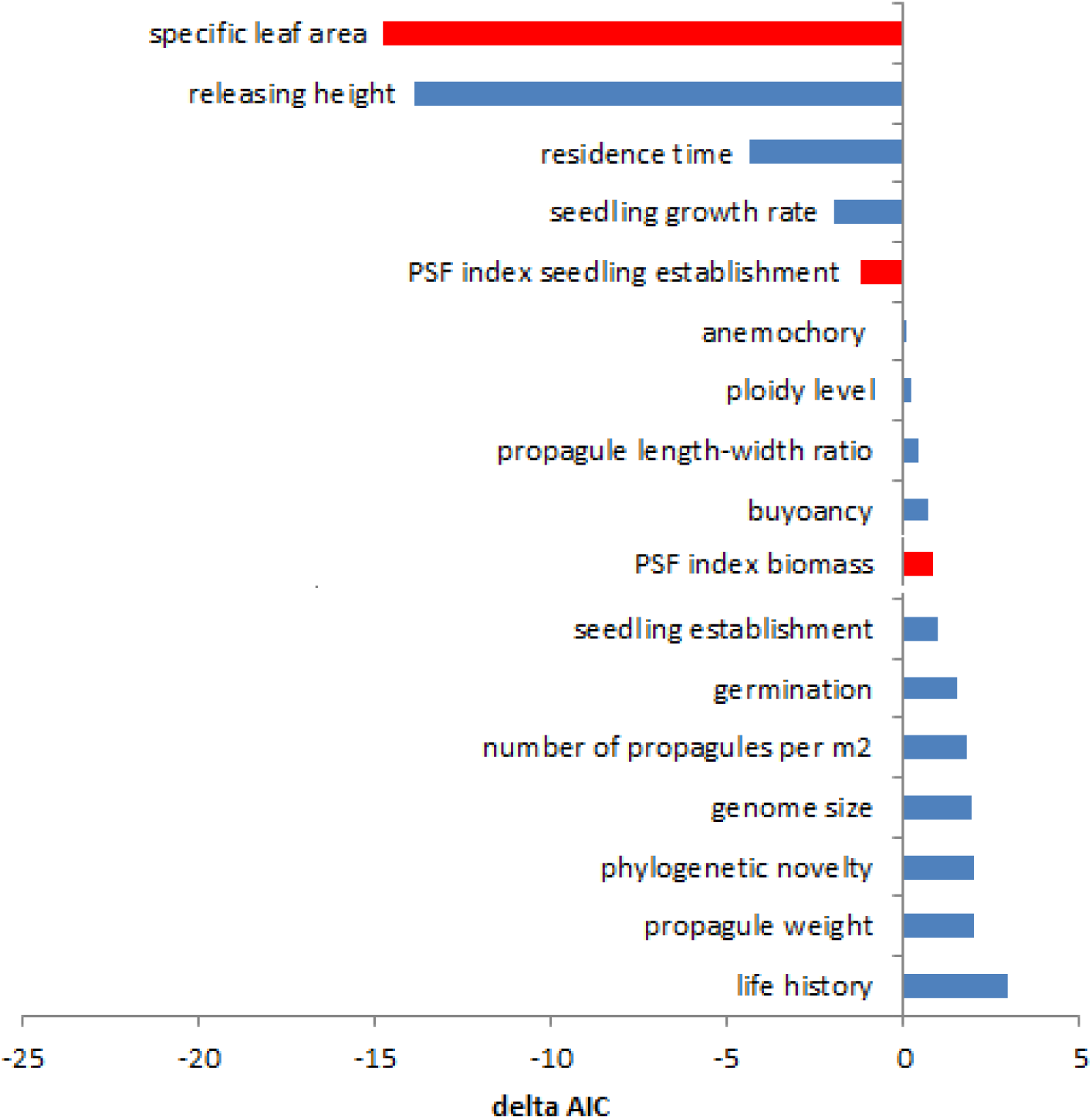
Delta AIC for models studying the effect of various species characteristics on invasive status. Negative values indicate significant contribution to explaining invasiveness. Red bars represent characteristics measured directly in our study, blue bars characteristics taken from other studies (see Methods).

### Statistical analyses

First, we tested for the effect of soil treatment (i.e. conditioned or control soil) on seedling establishment (here used as number of established seedlings divided by number of sown seeds, i.e. by 10) and on square-root transformed total biomass (sum of aboveground and belowground biomasses) in the feedback phase for each species separately. We used linear mixed effect models using the R-package ‘lmerTest’ (Kuznetsova et al., 2017), with population and pairs of pots as random effects. These tests provide information on existence of non-neutral intraspecific PSF in each species. For each species, we then calculated PSF index for seedling establishment and total biomass as ln(x/s) where x is performance of each individual plant when grown in the conditioned soil and s is performance of a plant grown in the paired control pot, as suggested by Brinkman et al. (2010). PSF index values for all studied species are presented in Table S2. An index value of less than zero indicates a negative feedback, meaning the plant performs worse in the conditioned soil than in the control soil, while a value greater than zero indicates a positive feedback, meaning the plant performs better in the conditioned soil than in the control soil.

The data from the feedback phase were further analyzed for all the species combined using linear mixed effect models with species, population and pairs of pots as random effects, and seedling establishment and square root transformed biomass from the feedback phase as dependent variables. Soil treatment, invasive status, MRT and their interactions were used as explanatory variables. The same tests were performed with species frequency and species cover in the field as alternative measures of plant invasiveness. The results of these analyses were consistent with those using invasive status and are presented in the Supporting information (Tables S4-S5, Figs S1-S3). Similarly, we performed tests with phylogenetic novelty (existence of any native species of the same genus in the Czech Republic) instead of MRT. Species invasive status is used as an explanatory variable in these tests, thought biologically invasive status is a function of PSF and not the other way around. Despite this, we used invasive status as the explanatory variable because such a test allowed us to study the interactions between invasive status and soil conditioning, MRT and phylogenetic novelty. It is, however, important to keep this in mind when interpreting the results.

To test for the relative importance of PSF for plant invasiveness in comparison with other species characteristics, we first calculated an average value of PSF index for seedling establishment and for biomass for each species across all populations. Then we used generalized linear models with binomial error distribution with invasive status as a dependent variable and a species characteristic affecting plant invasiveness as an explanatory variable. We performed the model separately for each studied species characteristic and compared delta AIC of all models with a null model.

All analyses were performed twice, once with phylogenetic correction and once without it. To account for phylogenetic signal in the data, we first extracted a phylogenetic tree for the 68 species in our study from Daphne (Durka and Michalski, 2012). Second, the phylogenetic distance matrix was decomposed into its eigenvectors using PCoA in the R-package ‘ape’ (Paradis et al., 2004). The first three eigenvectors, that together explained more than 50 % of the variability in the data, were included as covariables in the analyses in order to correct for phylogenetic autocorrelation. Since the analyses with and without the phylogenetic correction produced very similar results, we present only the results of the phylogenetically informed analyses. All analyses were performed using R 2.13.2 (R Development Core Team, 2014).

## Results

Significant positive PSF for seedling establishment was shown in four invasive and eight non-invasive species, significant negative PSF in two invasive and six non-invasive species. Significant positive PSF for biomass was shown in three invasive and five non-invasive species, significant negative PFS for three invasive and twelve non-invasive species (Table S2).

Overall, seedling establishment was significantly related to the soil treatment with plants performing better in the cultivated soil compared to the control soil (F = 11.9, P = 0.001). There was a significant interaction of soil treatment and invasive status (F = 3.53, P = 0.05), with seedlings of invasive species profiting more from growing in conditioned soil, i.e. having more positive PSF, than seedlings of non-invasive species (Fig. 1). Biomass was significantly affected by the soil treatment (F = 24.5, P < 0.001) with plants growing worse in the conditioned soil compared to the control soil, i.e. showing negative PSF. There was no significant difference in the response of invasive and non-invasive species to the soil conditioning for plant biomass.

**Fig. 1:**
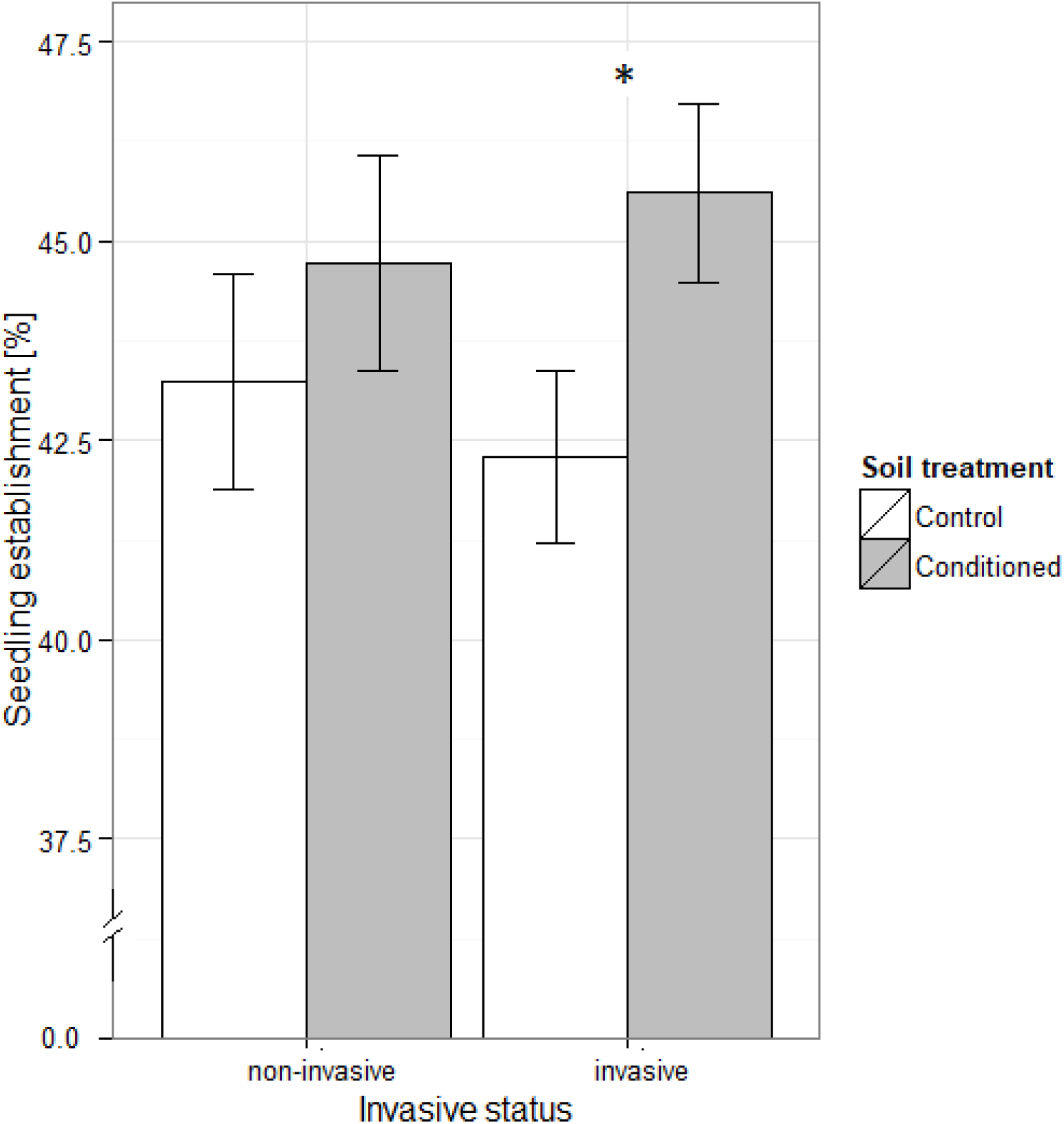
Seedling establishment of invasive and non-invasive species in control and conditioned soil. Asterisks indicate significant (P < 0.05) difference between control and conditioned soil. Better performance in conditioned soil compared to control indicates positive PSF, better performance in control soil compared to conditioned soil indicates negative PSF.

Neither seedling establishment nor biomass was significantly related to the interaction of MRT and soil treatment when all species were considered. However, triple interactions of MRT, soil treatment and invasion status were significant (F = 20.92, P < 0.001 for seedling establishment, F = 5.57, P = 0.018 for biomass). We therefore tested for the effect of MRT also separately for invasive and non-invasive species and found opposite relationships in the two categories (Fig. 2). While PSF of invasive species was positively correlated to MRT (F = 16.3, p < 0.001 for seedling establishment, Fig. 2a, F = 3.14, P = 0.076 for biomass, Fig. 2b), PSF of non-invasive species was negatively correlated to MRT (F = 4.76, P = 0.029 for seedling establishment, Fig. 2a, F = 7.28, P = 0.007 for biomass, Fig. 2b).

**Fig. 2:**
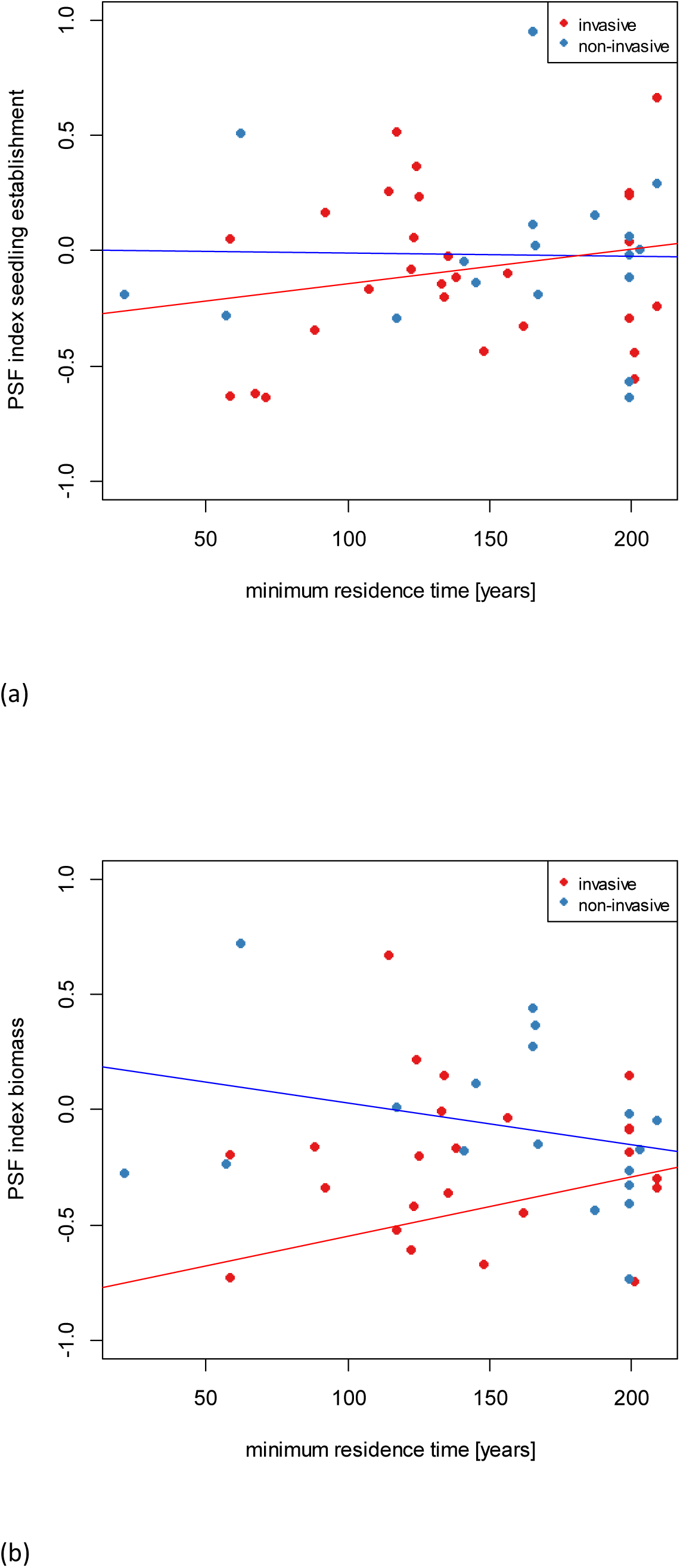
Dependence of PSF index for (a) seedling establishment and (b) biomass on minimum residence time for non-invasive and invasive plant species. Each data point represents mean PSF index of a single species.

Seedling establishment, but not biomass, was significantly affected by the interaction of the soil treatment and phylogenetic novelty of the species (F = 8.29, P = 0.004), while the interaction of phylogenetic novelty with invasive status and the triple interaction between the soil treatment, phylogenetic novelty and invasive status were not significant. Phylogenetically novel species, i.e. species that do not have a congener native to the country, had significantly better seedling establishment in the conditioned soil compared to the control soil, while species that do have a native congener performed comparably in the conditioned and the control soil (Fig. 3).

**Fig. 3:**
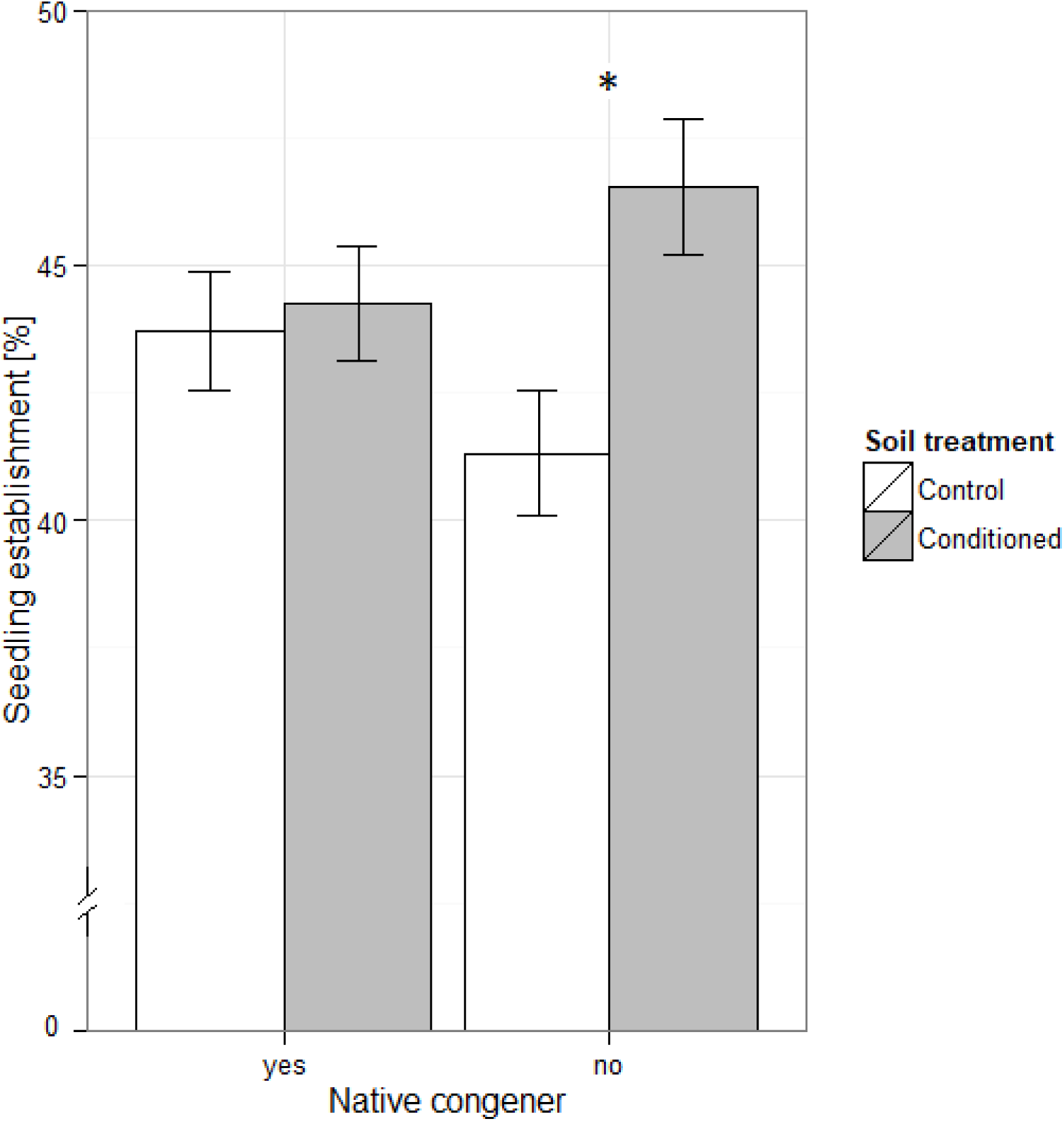
Seedling establishment of species that do and do not have a native congener in the Czech Republic in control and conditioned soil. Asterisks indicate significant (P < 0.05) difference between control and conditioned soil. Better performance in conditioned soil compared to control indicates positive PSF, better performance in control soil compared to conditioned soil indicates negative PSF.

The best predictors of invasive status were SLA, releasing height, MRT, seedling growth rate and PSF index for seedling establishment (Fig. 4). Similar results were obtained when predicting species frequency and maximum cover in the field (Table S5).

## Discussion

In the present study, we demonstrated on a large set of species that invasive species have more positive PSF for seedling establishment, but not for biomass, than alien non-invasive species, and we showed that PSF for seedling establishment belonged to the best predictors of plant invasiveness compared to a wide range of other species characteristics. We also showed that PSF is affected by phylogenetical novelty of the aliens and that it depends on residence time of the species.

Our study considered 68 alien plant species and 17 species characteristics, including PSF, providing a unique analysis of the relative importance of different predictors of plant invasiveness. The five most important predictors were specific leaf area, height, seedling growth rate, MRT and PSF for seedling establishment. We found that tall plants with high SLA and fast seedling growth rates are more likely to become invasive, which corresponds to previous studies (Pyšek and Richardson, 2007, van Kleunen et al., 2010, Ordonez et al., 2010). High specific leaf area is correlated with fast growth rate (Grotkopp et al., 2002) and height is positively associated with long-distance dispersal ability and with ability to compete for light (Thomson et al., 2011). Together, these traits allow plants to better disperse to and succeed in the disturbed habitats. We found that alien plants with longer residence time are more likely to be classified as invasive. This is also in line with other research (Pysek and Jarosik, 2005, Pysek et al., 2009), and our research highlights the importance of MRT in relation to many other invasion mechanisms, as has been previously done for woody plants (Pysek et al., 2009). Importantly, alien species are more likely to be classified as invasive if they have more positive PSF in their seedling establishment. PSF responses are not currently considered in programs that evaluate alien plants for invasiveness, and seedling establishment is not a commonly measured response variable in PSF research. Our results show that PSF especially for seedling establishment should receive more attention in future basic and applied studies since PSF was a better predictor of invasiveness than many other species characteristics commonly considered, such as size or shape of propagules, genome size or ploidy level.

We showed that invasive species have more positive PSFs for seedling establishment than non-invasive aliens, regardless of whether invasive success was categorical or based on maximum cover or frequency in the field. PSFs have been shown to allow invasive plants to gain dominance over native species in many previous studies (e.g., Kulmatiski et al., 2008, Meisner et al., 2014, van Grunsven et al., 2007), and to be positively linked with plant abundance in the field (Klironomos, 2002, Mangan et al., 2010, but see Reinhart, 2012, Maron et al., 2016 for no or opposite effect). Our study shows that PSFs also differentiate invasive from less successful alien species, even though the differences were not too prominent. There are many possible mechanisms that could explain this result. Successful alien plant species may be less regulated by enemies and/or benefit more from the interactions with soil mutualists (Fitter, 2005, Reinhart and Callaway, 2006, Menzel et al., 2017). In addition, successful aliens may have an intrinsic potential to improve the soil environment to their favor, for example via affecting decomposition rates and nutrient cycling (Ehrenfeld, 2004, Wolfe and Klironomos, 2005, Hu et al., 2018).

PSF for seedling establishment played more important role for plant invasiveness than PSF for biomass in our study system, confirming that early stages of plant lives are crucial for successful invasion (Gioria and Pysek, 2017). This is an important finding because seedling establishment is not commonly considered in PSF research (Kardol et al., 2013) and we know little about how plant-soil interactions affect seedling recruitment. A few studies have shown that PSF of seedlings may largely differ from that of adults and tends to be more positive (Dudenhoffer et al., 2018, Florianova and Munzbergova, 2018). This may be because seedlings have less developed root system and can thus benefit more from associations with mycorrhizal fungi (Aldrich-Wolfe 2007; van der Heijden 2004). On the other hand, seedlings may be very vulnerable to negative effects of soil pathogens in some systems (Packer and Clay, 2000, Liu et al., 2015). Number of established seedlings in our study could have also been affected by different germination rates that might be subject to PSF as well. Some plants release specific root exudates that stimulate plant growth-promoting rhizobacteria (van Loon, 2007, Vacheron et al., 2013, Hu et al., 2018), and there is some evidence, mostly from agricultural systems, that these can positively affect seed germination (e.g., Kloepper et al., 1991, Wu et al., 2016). Some plants have also been shown to release chemicals that directly stimulate germination of seeds. This has mostly been observed in interspecific interactions and the focal seeds are typically of parasitic plants (Netzly et al., 1988, Hameed et al., 1973, Fernandez-Aparicio et al., 2009, Fernandez-Aparicio et al., 2008). However, it is possible that similar mechanisms are involved in intraspecific PSF.

PSFs of aliens are expected to become more negative over time due to adaptation and accumulation of pathogens (Dostal et al., 2013, Flory and Clay, 2013), but this pattern has received mixed support (Diez et al., 2010, Speek et al., 2015, McGinn et al., 2018). Our study suggests that this inconsistency could be due to the lack of explicit consideration of invasive status. In our study, non-invasive aliens showed the expected negative pattern with MRT, while invasive aliens showed an opposite relationship. Most studies examining the release from belowground enemies have focused on problematic invaders and little is known about the release of non-invasive aliens (Callaway et al., 2004, Reinhart et al., 2003, van Grunsven et al., 2007, Gundale et al., 2014, Maron et al., 2014, Yang et al., 2013). Alien plants might differ in the number of enemy taxa that initially accompany them to the new regions (MacLeod et al., 2010) and in the rate at which they subsequently accumulate enemies (Dickie et al., 2017). However, plant-soil mutualist interactions may also accumulate over time (Dickie et al., 2017) and the balance between accumulation of pathogens and mutualists will determine the net PSF pattern. It has been suggested that mutualist limitation, even though not so common and often temporary, may reduce the rate of population growth and abundance and prevent a naturalized species from becoming invasive (Dickie et al., 2017). Moreover, it has been documented that problematic invasive plants may benefit from novel soil mutualists (Reinhart and Callaway, 2004, Reinhart and Callaway, 2006, Sun and He, 2010), and more studies are needed to determine if such patterns differ between invasive and non-invasive aliens.

Alien species that are phylogenetically related to the native flora should be more likely to share species-specific pathogens (Bufford et al., 2016, Vacher et al., 2010, Parker et al., 2015, Gilbert and Parker, 2016). They should therefore develop more negative PSFs than phylogenetically novel species that are only attacked by generalist pathogens, having usually considerably smaller effect on plants than specialists (Colautti et al., 2004). On the contrary, soil mutualists, such as AM fungi, whose accumulation in the soil might lead to positive PSF, usually have low level of endemism (Davison et al., 2015) and lower host specificity (Smith and Read, 2008, Molina and Horton, 2015), and the alien species can thus usually benefit from their presence regardless of their phylogenetic relatedness to native flora (Yang et al., 2014, Richardson et al., 2000, McGinn et al., 2016). Previous studies have demonstrated that plants grow poorly when grown in soil conditioned by a close relative (Callaway et al., 2013, Dostal and Paleckova, 2011, Kempel et al., 2018, but see Kutakova et al., 2018, Munzbergova and Surinova, 2015, Mehrabi and Tuck, 2015). However, our study aimed to examine whether or not novelty of aliens can explain PSF patterns across many alien plant species, all growing in resident soil. Indeed, we found the expected pattern that alien species with a native congener in the Czech Republic have more negative PSFs than phylogenetically novel species. Our results together with other literature show that the presence of closely related native plant species has negative effects on alien species via the activity of herbivores, phytophagous insects, above-ground or below-ground pathogens (e.g., Connor et al., 1980, Flory and Clay, 2013, Hill and Kotanen, 2009, Parker et al., 2015).

The majority of species in our study showed neutral PSF, which contradicts the general worldwide pattern that most species show negative intraspecific PSF (e.g., Kulmatiski et al., 2008, Petermann et al., 2008). This is likely to be caused by our study focusing only on aliens, which may be at least partly released from negative effects of their host-specific enemies (see also the neutral and positive PSFs of invasives reported by Callaway et al., 2004, Reinhart and Callaway, 2006, Meisner et al., 2014, Suding et al., 2013, Engelkes et al., 2008, Levine et al., 2006). In addition, the strength of PSFs might depend on duration of soil conditioning or type of soil used (van de Voorde et al., 2012, Lepinay et al., 2018). In our study, we used local common garden compost mixed with sand while the majority of previous studies used soil collected directly in the localities. Our soil could thus initially contain lower abundances of species specific pathogens, which would cause slower pathogen accumulation during the conditioning phase causing experimental results to be less negative. The soil also contained more nutrients reducing the likelihood of negative PSF due to nutrient depletion. As the initial soil used for the experiment likely has significant effects on the results of PSF (van de Voorde et al., 2012, Png et al., 2018, Gundale et al., 2019), it would be better to use more soil types in our experiment. Doing this would, however, make our experiment unfeasible. In addition, other previous studies demonstrated that the effects of specific soil used are much weaker than the effects of soil cultivation and do not interact (Hemrova et al., 2016).

Our results have implications for future studies in other regions, for future research on PSFs, and for biosecurity screening of alien plants. Our research can serve as a guide for distributed observations and experiments in other regions of the world. By considering multiple species characteristics that might explain invasiveness and using consistent methodology across all aliens in the PSF experimentation, our methods allow the importance of different species characteristics to be ranked and the magnitude and direction of PSFs to be compared across species. If such an approach was replicated in other regions, this would enable more sophisticated meta-analyses of invasion mechanisms. Our research highlights the need for future work on PSFs, particularly focusing on the mechanisms by which these feedbacks influence seedling establishment. Finally, biosecurity screening processes for importing new aliens, such as the Australian Weed Risk Assessment System (Gordon et al., 2008), should consider evaluating aliens for positive PSFs in case future studies in other regions confirm our results. Such experimentation is relatively fast and inexpensive to implement and PSF was found to contribute to predicting invasiveness in our system.

## Acknowledgements

We would like to thank T. Knight for assistance with writing the manuscript, L. Moravcová for providing information on localities of our studied species in the Czech Republic and their germination requirements, K. Skácelová for help with seed collection and I. Jarošincová, I. Chmelařová, D. Cmíral and M. Chmelař for help with maintaining the experiment. The study was supported by Czech Science Foundation (project GAČR 16-09659S). It was also partly supported by RVO 67985939 and Czech Ministry of Education, Youth and Sports (MŠMT).

## Author contributions

AA and ZM designed the study, AA and PK collected data, AA and ZM performed the statistical analyses and wrote the manuscript.

