## Supplemental information for "Plant-soil feedback contributes to predicting plant invasiveness of 68 alien plant species differing in invasive status"

Table S1: Summary characteristics of the neophytes in the Czech flora studied in the present paper, shown separately for invasive (n = 27) and non-invasive (n = 41) alien species. Differences between invasive and non-invasive species tested with chi-square test or one-way ANOVA, for minimum residence time and species frequency on square-root and ln transformed data, respectively.

| | Non-invasive | Invasive | Difference |
|---|---|---|---|
| Life history | Annuals 15 (37 %)<br>Monocarpic perennials 9 (22 %)<br>Polycarpic perennials 17 (41 %) | Annuals 13 (48 %)<br>Monocarpic perennials 4 (15 %)<br>Polycarpic perennials 10 (37 %) | $X^2$ = 1.378; df = 2; P = 0.502 |
| Minimum residence time (years, mean ± S.D.) | 144.7 ± 54.6 | 156.5 ± 37.6 | F = 1.289; df = 1, 59; P = 0.261 |
| Max cover (%, mean ± S.D.<br>(min, max) | 35.6 ± 31.4<br>(1-88) | 70.4 ± 35.2<br>(1-99) | F = 5.1427; df = 1, 55; P = 0.027 |
| Species frequency (number of occupied grids, mean ± S.D.)<br>(min, max) | 392.3 ± 534.8<br>(5-1851) | 657.3 ± 563.5<br>(30-2190) | F = 7.722; df = 1, 65; P = 0.007 |
| Phylogenetic novelty<br>(absence of native congener) | Yes – 16 (40 %)<br>No – 25 (60 %) | Yes – 13 (48 %)<br>No – 14 (52 %) | $X^2$ = 0.591; df = 1; P = 0.442 |

Table S2: List of studied species, their description and PSF indices for biomass and seedling establishment. P-values indicate whether seedling establishment or biomass in the feedback phase are affected by soil treatment (conditioned, not-conditioned). Significant values ($P < 0.05$) are in bold, marginally significant values ($P < 0.1$) in italics. Life history: an – annual, mono – monocarpic perennial, per – polycarpic perennial. Invasion status: inv – invasive, non-inv – non-invasive. Minimum residence time – number of years elapsed since the first record of occurrence in the Czech Republic. Species frequency – number of colonized quadrants of basic cells in grid mapping. Maximum cover – maximum cover in the field. Number of populations – number of populations studied. NA – information not available.

| Species | Family | Life history | Invasion status | Minimum residence time [years] | Species occurrence | Maximum cover [%] | Seedling establishment | | Biomass | | Number of populations |
|---|---|---|---|---|---|---|---|---|---|---|---|
| | | | | | | | PSF index (mean ± sd) | p-value | PSF index (mean ± sd) | p-value | |
| Abutilon theophrasti Med. | Malvaceae | an | non-inv | 124 | 61 | 4 | **0.372 ± 0.247** | **0.014** | 0.018 ± 0.537 | 0.649 | 3 |
| Amaranthus albus L. | Amaranthaceae | an | non-inv | 125 | 181 | 63 | 0.24 ± 0.523 | 0.251 | -0.331 ± 0.817 | 0.485 | 3 |
| Amaranthus powellii S. Watson | Amaranthaceae | an | inv | 165 | 433 | 88 | *0.06 ± 0.627* | *0.063* | -0.075 ± 0.47 | 0.866 | 2 |
| Amaranthus retroflexus L. | Amaranthaceae | an | inv | 200 | 931 | 63 | **0.686 ± 0.881** | **0.001** | -0.093 ± 0.763 | 0.168 | 3 |
| Ambrosia artemisiifolia L. | Asteraceae | an | inv | 135 | 92 | 1 | -0.022 ± 0.268 | 0.588 | 0.07 ± 1.263 | 0.761 | 4 |
| Ambrosia trifida L. | Asteraceae | an | non-inv | 58 | 8 | 88 | 0.055 ± 0.58 | 0.111 | **-0.183 ± 0.337** | **0.013** | 3 |
| Antirrhinum majus L. | Plantaginaceae | mono | non-inv | 199 | 70 | 3 | **0.245 ± 0.648** | **0.034** | -0.019 ± 0.613 | 0.512 | 3 |
| Asclepias syriaca L. | Apocynaceae | per | inv | 117 | 86 | 88 | *0.161 ± 0.55* | *0.075* | **0.407 ± 0.497** | **0.008** | 3 |
| Aster lanceolatus Willd. | Asteraceae | per | inv | NA | 189 | NA | **-0.747 ± 0.358** | **<0.001** | **0.439 ± 0.464** | **0.005** | 3 |
| Bidens frondosus L. | Asteraceae | an | inv | 124 | 1360 | 88 | -0.012 ± 0.57 | 0.836 | 0.162 ± 0.343 | 0.393 | 2 |
| Bunias orientalis L. | Brassicaceae | mono | inv | 162 | 344 | 88 | *-0.324 ± 0.5* | *0.053* | *-0.441 ± 0.675* | *0.059* | 2 |
| Cannabis ruderalis Janisch. | Cannabinaceae | an | inv | 150 | 30 | 3 | 0.01 ± 0.294 | 0.298 | 0.157 ± 0.861 | 0.304 | 2 |
| Cardamine chelidonia L. | Brassicaceae | mono | non-inv | 88 | 20 | NA | -0.34 ± 0.538 | 0.2 | -0.063 ± 0.739 | 0.509 | 2 |
| Claytonia alsinoides Sims | Montiaceae | an | non-inv | 67 | NA | NA | **-0.618 ± 0.585** | **0.003** | **-1.772 ± 1.243** | **<0.001** | 2 |
| Collomia grandiflora Lindl. | Polemoniaceae | an | non-inv | 138 | 5 | NA | -0.111 ± 0.583 | 0.902 | -0.135 ± 0.514 | 0.268 | 3 |
| Conyza canadensis (L.) Cronq. | Asteraceae | an | inv | 268 | 1463 | 88 | **0.957 ± 1.005** | **0.025** | 0.475 ± 0.656 | 0.133 | 3 |
| Datura stramonium L. | Solaginaceae | an | non-inv | 209 | 286 | 38 | **0.586 ± 1.013** | **<0.001** | 0.342 ± 1.913 | 0.683 | 4 |
| Digitalis purpurea L. | Plantaginaceae | mono | non-inv | 228 | 650 | 63 | -0.375 ± 0.858 | 0.879 | -0.079 ± 0.733 | 0.279 | 2 |
| Duchesnea indica (Andrew) Focke | Rosaceae | per | non-inv | 58 | 38 | 1 | **-0.626 ± 0.501** | **0.009** | **-0.593 ± 1.26** | **0.015** | 2 |
| Echinocystis lobata (Michx.) Torr. et Gray | Cucurbitaceae | an | inv | 107 | 253 | 88 | -0.166 ± 0.366 | 0.803 | **-1.26 ± 0.82** | **0.027** | 1 |
| Echinops sphaerocephalus L. | Asteraceae | per | inv | 147 | 729 | 88 | -0.156 ± 0.45 | 0.727 | 0.009 ± 0.289 | 0.684 | 2 |
| Epilobium ciliatum Rafin. | Onagraceae | per | inv | 92 | 1851 | 38 | **0.168 ± 0.784** | **0.011** | -0.176 ± 1.454 | 0.831 | 4 |
| Erigeron annuus (L.) Pers. | Asteraceae | mono | inv | 134 | 388 | 63 | 0.110 ± 0.171 | 0.341 | 0.414 ± 0.607 | 0.104 | 1 |
| Galinsoga parviflora Cav. | Asteraceae | an | inv | 138 | 1198 | 88 | *-0.17 ± 0.303* | *0.1* | 0.248 ± 1.07 | 0.314 | 4 |
| Galinsoga quadriradiata Ruiz et Pavón | Asteraceae | an | inv | 117 | 1215 | 88 | **0.52 ± 0.436** | **0.007** | -0.113 ± 1.244 | 0.591 | 1 |
| Geraniaceaenium pyrenaicum Burm. fil. | Geraniaceae | per | non-inv | 199 | 530 | 38 | 0.125 ± 0.39 | 0.139 | -0.101 ± 0.285 | 0.52 | 3 |
| Helianthus tuberosus L. | Asteraceae | per | inv | 133 | 578 | 99 | -0.139 ± 0.619 | 0.859 | 0.081 ± 1.602 | 0.177 | 2 |
| Heracleum mantegazzianum Sommier et Levier | Apiaceae | mono | inv | 156 | 694 | 99 | -0.097 ± 0.18 | 0.203 | -0.028 ± 0.526 | 0.565 | 2 |
| Hesperis matronalis L. | Brassicaceae | per | non-inv | 201 | 552 | 18 | **-0.482 ± 0.502** | **0.007** | **-0.469 ± 0.612** | **0.012** | 2 |
| Chenopodium pumilio R. Br. | Amaranthaceae | an | non-inv | 128 | 113 | 3 | **-0.442 ± 0.365** | **0.015** | **-1.719 ± 1.449** | **0.001** | 1 |
| Chenopodium strictum Roth | Amaranthaceae | an | non-inv | NA | 532 | 2 | **-0.611 ± 0.697** | **0.047** | **-0.321 ± 0.646** | **0.014** | 5 |
| Impatiens glandulifera Royle | Balsaminaceae | an | inv | 122 | 1214 | 90 | -0.079 ± 0.494 | 0.92 | **-0.471 ± 0.633** | **0.005** | 3 |

| Species | Family | Life history | Invasion status | Minimum residence time [years] | Species occurrence | Maximum cover [%] | Seedling establishment | | Biomass | | Number of populations |
|---|---|---|---|---|---|---|---|---|---|---|---|
| | | | | | | | PSF index (mean ± sd) | p-value | PSF index (mean ± sd) | p-value | |
| Impatiens parviflora DC. | Balsaminaceae | an | inv | 148 | 2190 | 99 | -0.434 ± 0.335 | 0.645 | *-0.442 ± 0.642* | *0.093* | 2 |
| Imperatoria ostruthium L. | Apiaceae | per | non-inv | 209 | 123 | 63 | *-0.239 ± 0.299* | *0.077* | 0.068 ± 1.102 | 0.863 | 3 |
| Iva xanthiifolia Nutt. | Asteraceae | an | non-inv | 71 | 55 | 88 | -0.637 ± 0.891 | 0.35 | **-1.331 ± 1.168** | **0.017** | 1 |
| Kochia scoparia (L.) Schrader | Amaranthaceae | an | inv | 199 | 138 | 88 | -0.288 ± 0.55 | 0.133 | -0.046 ± 0.279 | 0.435 | 3 |
| Lepidium densiflorum Schrader | Brassicaceae | mono | non-inv | 114 | 244 | 38 | **0.259 ± 0.281** | **0.046** | **0.747 ± 0.645** | **0.008** | 1 |
| Lupinus polyphyllus Lindl. | Fabaceae | per | inv | 123 | 1167 | 70 | -0.092 ± 0.374 | 0.682 | -0.142 ± 0.572 | 0.692 | 3 |
| Lychnis coronaria (L.) Desr. | Caryophyllaceae | mono | non-inv | 139 | 67 | 2 | -0.521 ± 0.636 | 0.731 | -0.454 ± 0.469 | **0.015** | 3 |
| Lysimachia punctata. L. | Primulaceae | per | non-inv | 199 | 423 | 2 | *0.254 ± 0.575* | *0.073* | 0.283 ± 0.875 | 0.294 | 2 |
| Matricaria discoidea DC. | Asteraceae | an | non-inv | 165 | 1733 | 88 | **0.954 ± 0.548** | **<0.001** | 0.393 ± 1.459 | 0.101 | 2 |
| Medicago sativa L. | Fabaceae | per | non-inv | 199 | 1020 | 63 | *-0.113 ± 0.403* | *0.078* | -0.183 ± 0.485 | 0.246 | 2 |
| Mimulus guttatus DC. | Phrymaceae | per | non-inv | 165 | 163 | 13 | 0.096 ± 0.719 | 0.104 | **0.33 ± 0.58** | **<0.001** | 4 |
| Oenothera biennis L. | Onagraceae | mono | non-inv | 187 | 621 | NA | *-0.132 ± 0.501* | *0.078* | -0.237 ± 0.511 | 0.114 | 3 |
| Oenothera glazioviana M. Micheli | Onagraceae | mono | non-inv | 128 | 177 | NA | 0.183 ± 0.282 | 0.183 | **-0.146 ± 0.291** | **0.017** | 2 |
| Oxalis dillenii Jacq. | Oxalidaceae | mono | inv | NA | 75 | 2 | 0.201 ± 0.454 | 0.404 | **0.463 ± 0.966** | **0.018** | 3 |
| Oxalis fontana Bunge | Oxalidaceae | mono | non-inv | 166 | 1043 | 38 | 0.024 ± 0.535 | 0.458 | **0.643 ± 1.127** | **0.048** | 1 |
| Phytolacca esculenta Van Houtte | Phytolaccaceae | per | non-inv | 62 | 35 | NA | **0.51 ± 0.871** | **<0.001** | **0.916 ± 2.127** | **<0.001** | 3 |
| Rudbeckia hirta L. | Asteraceae | per | non-inv | 145 | 77 | 38 | 0.068 ± 0.556 | 0.208 | 0.193 ± 0.556 | 0.317 | 2 |
| Rudbeckia laciniata L. | Asteraceae | per | inv | 159 | 343 | 13 | 0.217 ± 0.477 | 0.118 | -0.013 ± 0.983 | 0.846 | 1 |
| Rumex alpinus L. | Polygonaceae | per | non-inv | 199 | 65 | 90 | -0.026 ± 0.305 | 0.865 | **-0.405 ± 0.429** | **<0.001** | 3 |
| Rumex longifolius DC. | Polygonaceae | per | non-inv | NA | 46 | NA | 0.064 ± 0.38 | 0.423 | -0.203 ± 0.623 | 0.178 | 2 |
| Rumex patientia L. subsp. patientia | Polygonaceae | per | inv | 157 | 32 | 38 | -0.111 ± 0.352 | 0.777 | -0.172 ± 0.442 | 0.402 | 2 |
| Rumex thyrsiflorus Fingerh. | Polygonaceae | per | non-inv | NA | 401 | 13 | **-0.765 ± 0.76** | **0.031** | *-0.206 ± 0.526* | *0.079* | 3 |
| Scutellaria altissima L. | Lamiaceae | per | non-inv | 117 | 18 | NA | -0.29 ± 0.521 | 0.395 | 0.037 ± 0.218 | 0.705 | 1 |
| Sedum hispanicum L. | Crassulaceae | per | non-inv | NA | 334 | 2 | **0.441 ± 0.573** | **0.003** | -0.573 ± 1.357 | 0.278 | 3 |
| Sedum rupestre L. subsp. erectum t´Hart | Crassulaceae | per | non-inv | NA | 61 | 38 | 0.091 ± 0.474 | 0.342 | **-1.779 ± 1.147** | **<0.001** | 1 |
| Senecio inaequidens DC. | Asteraceae | per | non-inv | 21 | 116 | NA | -0.186 ± 0.801 | 0.939 | 0.009 ± 1.281 | 0.512 | 3 |
| Setaria faberi F. Herrmann | Poaceae | an | non-inv | 57 | 19 | 1 | -0.28 ± 0.608 | 0.4 | *-0.211 ± 0.321* | *0.07* | 1 |
| Sisymbrium altissimum L. | Brassicaceae | an | non-inv | 203 | 287 | 88 | 0.008 ± 0.588 | 0.9 | **0.633 ± 1.72** | **0.034** | 4 |
| Sisymbrium loeselii L. | Brassicaceae | an | inv | 199 | 409 | 88 | **-0.668 ± 1.005** | **0.002** | **-0.37 ± 0.996** | **0.009** | 3 |
| Sisymbrium strictissimum L. | Brassicaceae | per | non-inv | 199 | 325 | 13 | -1.226 ± 0.661 | 0.133 | **-0.584 ± 1.654** | **<0.001** | 1 |
| Solidago canadensis | Asteraceae | per | inv | 180 | 1202 | 99 | 1.644 ± 0.423 | 0.389 | 0.097 ± 0.498 | 0.392 | 1 |
| Telekia speciosa (Schreber) Baumg. | Asteraceae | per | inv | 198 | 305 | 1 | -0.074 ± 0.659 | 0.813 | -0.066 ± 0.889 | 0.776 | 3 |
| Trifolium hybridum L. | Fabaceae | mono | non-inv | 199 | 1783 | 88 | 0.031 ± 0.497 | 0.353 | *0.361 ± 1.342* | *0.093* | 4 |
| Veronica persica Poiret | Plantaginaceae | an | non-inv | 209 | 1838 | 63 | **0.296 ± 0.471** | **0.002** | -0.018 ± 0.348 | 0.707 | 3 |
| Vicia grandiflora Scop. | Fabaceae | an | non-inv | 141 | 42 | NA | -0.041 ± 0.325 | 0.922 | *-0.120 ± 0.367* | *0.085* | 3 |
| Xanthium albinum (Widd.) H. Scholtz et Sukopp | Asteraceae | an | non-inv | 167 | 106 | 2 | -0.142 ± 0.303 | 0.443 | -0.048 ± 0.372 | 0.593 | 3 |

Table S3: Abiotic characteristics of the used soil prior to the conditioning phase. Values show the mean and stardard deviation of six samples. The analyses were performed by the Analytical Laboratory of Institute of Botany, Czech Academy of Sciences, Průhonice. The methods used for the analyses are described in detail in Raabova et al. (2008).

| | mean ± sd |
|---|---|
| pH($H_2$0) | 7.77 ± 0.03 |
| pH(KCl) | 7.70 ± 0.01 |
| total N [%] | 0.07 ± 0.02 |
| total C [%] | 1.01 ± 0.20 |
| exchangeable Ca [mg/kg] | 1259.35 ± 125.63 |
| exchangeable Mg [mg/kg] | 105.03 ± 11.53 |
| exchangeable K [mg/kg] | 278.60 ± 21.61 |
| exchangeable P [mg/kg] | 39.99 ± 1.12 |
| total P [mg/kg] | 159.46 ± 46.70 |

Table S4: Results of mixed effect models with i) seedling establishment and ii) biomass from the feedback phase as dependent variables and phylogenetic eigenvectors (axes 1-3), MRT, soil treatment, measure of invasion and their interaction as explanatory variables. Measure of invasion: invasion status, square-root transformed species frequency and maximum cover, respectively. Species, populations and pairs of pots were used as random effects. DenDF - Satterthwaite approximation for degrees of freedom. NumDF = 1 for all variables. Significant values (P < 0.05) are in bold, marginally significant values (P < 0.1) are in italics. Species frequency – number of colonized quadrants of basic cells in grid mapping. Maximum cover – based on maximum cover in the field, taken from Pladias, transformed into a discrete variable with three levels (low cover < 10 %, medium 11-50 %, high > 50 %).

| | | | axis1 | axis2 | axis3 | MRT | soil treatment | measure of invasion | meas. of invasion * soil treat. |
|---|---|---|---|---|---|---|---|---|---|
| Seedling establishment | invasion status | DenDF | 161.3 | *160.2* | 160.4 | **161.0** | **3037.1** | 160.6 | *3037.0* |
| | | F | 1.54 | *3.37* | 0.00 | **8.26** | **11.90** | 0.05 | *3.53* |
| | | P | 0.216 | *0.068* | 0.973 | **0.005** | **0.001** | 0.823 | *0.050* |
| | species frequency | DenDF | 159.4 | *158.2* | 158.6 | **158.6** | 2996.4 | **330.0** | **2999.4** |
| | | F | 1.73 | *3.36* | 0.07 | **8.34** | 1.27 | **5.36** | **9.88** |
| | | P | 0.190 | *0.069* | 0.259 | **0.004** | 0.259 | **0.005** | **0.002** |
| | maximum cover | DenDF | 134.3 | **133.2** | 133.5 | **133.9** | **2546.7** | *133.6* | **2546.4** |
| | | F | 0.00 | **6.92** | 0.11 | **4.71** | **10.78** | *2.74* | **5.61** |
| | | P | 0.965 | **0.010** | 0.739 | **0.032** | **0.001** | *0.069* | **0.004** |
| Biomass | invasion status | DenDF | 161.3 | 160.7 | *160.8* | 161.1 | **3026.7** | 160.9 | 3026.5 |
| | | F | 1.45 | 0.01 | *2.98* | 1.83 | **24.50** | 0.47 | 1.50 |
| | | P | 0.230 | 0.920 | *0.086* | 0.178 | **<0.001** | 0.494 | 0.220 |
| | species frequency | DenDF | 159.3 | 158.7 | 158.9 | 158.9 | **2987.2** | 312.5 | 2988.6 |
| | | F | 1.42 | 0.26 | 1.05 | 0.85 | **10.45** | 1.17 | 0.17 |
| | | P | 0.236 | 0.611 | 0.306 | 0.359 | **0.001** | 0.217 | 0.679 |
| | maximum cover | DenDF | 134.2 | 133.7 | 133.8 | 134.1 | **2545.5** | 133.9 | *2545.4* |
| | | F | 0.80 | 0.61 | 0.06 | 0.03 | **14.23** | 0.93 | *2.54* |
| | | P | 0.372 | 0.438 | 0.814 | 0.860 | **<0.001** | 0.396 | *0.079* |

Table S5: Comparison of delta AIC for models studying the effect of various species characteristics on invasive status, species frequency and maximum cover in the field. Underlined variables are data obtained in this study. For invasive status, we used generalized linear models with binomial error distribution, for species frequency linear models on square-root transformed data and for maximum cover linear models with multinomial error distribution ('multinom' function in 'nnet' package in R (Venables and Ripley, 2002)).

| | invasion status | | species frequency | | maximum cover | |
|---|---|---|---|---|---|---|
| | delta AIC | rank | delta AIC | rank | delta AIC | rank |
| specific leaf area | **-23.217** | **1** | **-170.788** | **1** | **-41.780** | **1** |
| releasing height | **-21.801** | **2** | **-92.106** | **2** | **-18.587** | **2** |
| MRT | **-6.814** | **3** | **-57.521** | **3** | **-11.483** | **3** |
| seedling growth rate | **-3.111** | **4** | **-26.486** | **4** | **-1.396** | **7** |
| PSF seedling establishment | **-1.871** | **5** | **-16.757** | **5** | **-0.024** | **11** |
| anemochory | 0.065 | 6 | **-14.767** | **6** | **-2.460** | **6** |
| ploidy level | 0.227 | 7 | **-6.385** | **8** | **-0.249** | **10** |
| propagule length-width ratio | 0.404 | 8 | **-8.217** | **7** | **-0.608** | **8** |
| buyoancy | 0.667 | 9 | 1.606 | 15 | 3.392 | 15 |
| PSF biomass | 0.814 | 10 | 0.997 | 10 | 3.508 | 16 |
| seedling establishment | 0.956 | 11 | 1.302 | 14 | 2.256 | 13 |
| germination | 1.519 | 12 | 1.284 | 13 | 2.685 | 14 |
| number of propagules per $m^2$ | 1.783 | 13 | 1.781 | 16 | 3.959 | 17 |
| genome size | 1.932 | 14 | 1.113 | 11 | **-4.354** | **5** |
| native congener | 1.982 | 15 | **-3.193** | **9** | 0.2077 | 12 |
| propagule weight | 1.991 | 16 | 1.245 | 12 | **-0.354** | **9** |
| life history | 2.979 | 17 | 3.012 | 17 | **-4.447** | **4** |

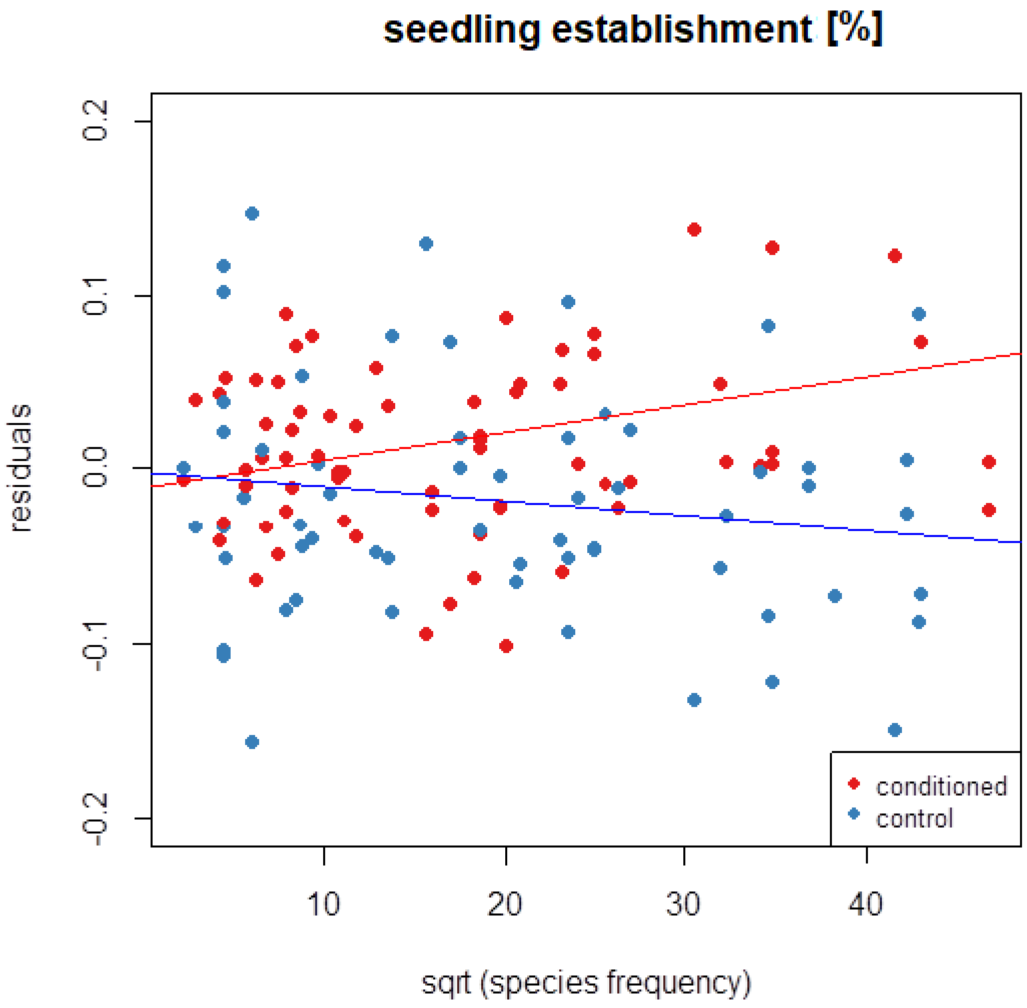


Fig. S1: Dependence of mean residuals for seedling establishment across species, after accounting for phylogenetic information, MRT and random effect of species, population and pairs of pots, on square-root transformed species frequency in quadrants of the basic grid mapping cells for conditioned and control soil. Better performance in conditioned soil compared to control indicates positive PSF, better performance in control soil compared to conditioned soil indicates negative PSF.

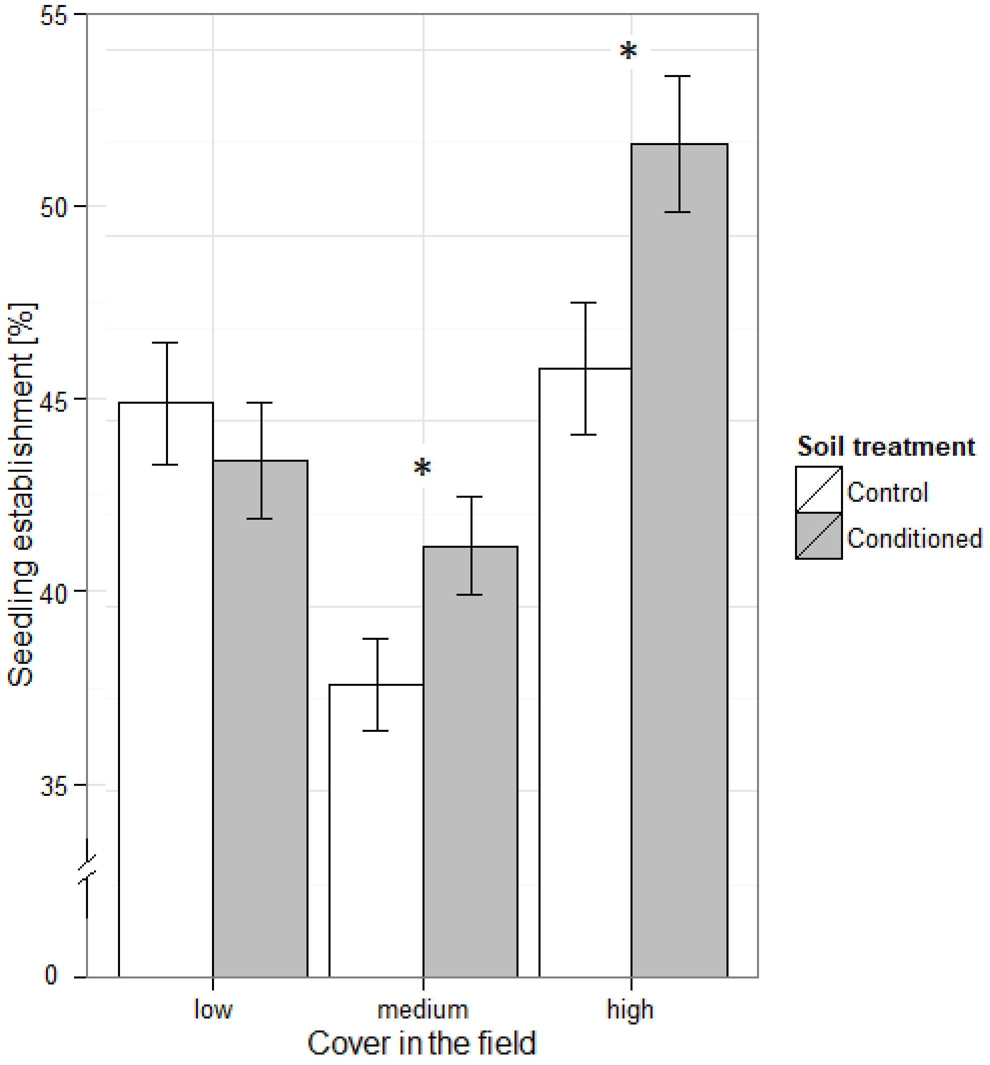


Fig. S2: Seedling establishment of species with low (< 10 %), medium (11-50 %) and high (> 50 %) maximum cover in the field in control and conditioned soil. Asterisks indicate significant (P < 0.05) difference between control and conditioned soil. Better performance in conditioned soil compared to control indicates positive PSF, better performance in control soil compared to conditioned soil indicates negative PSF.

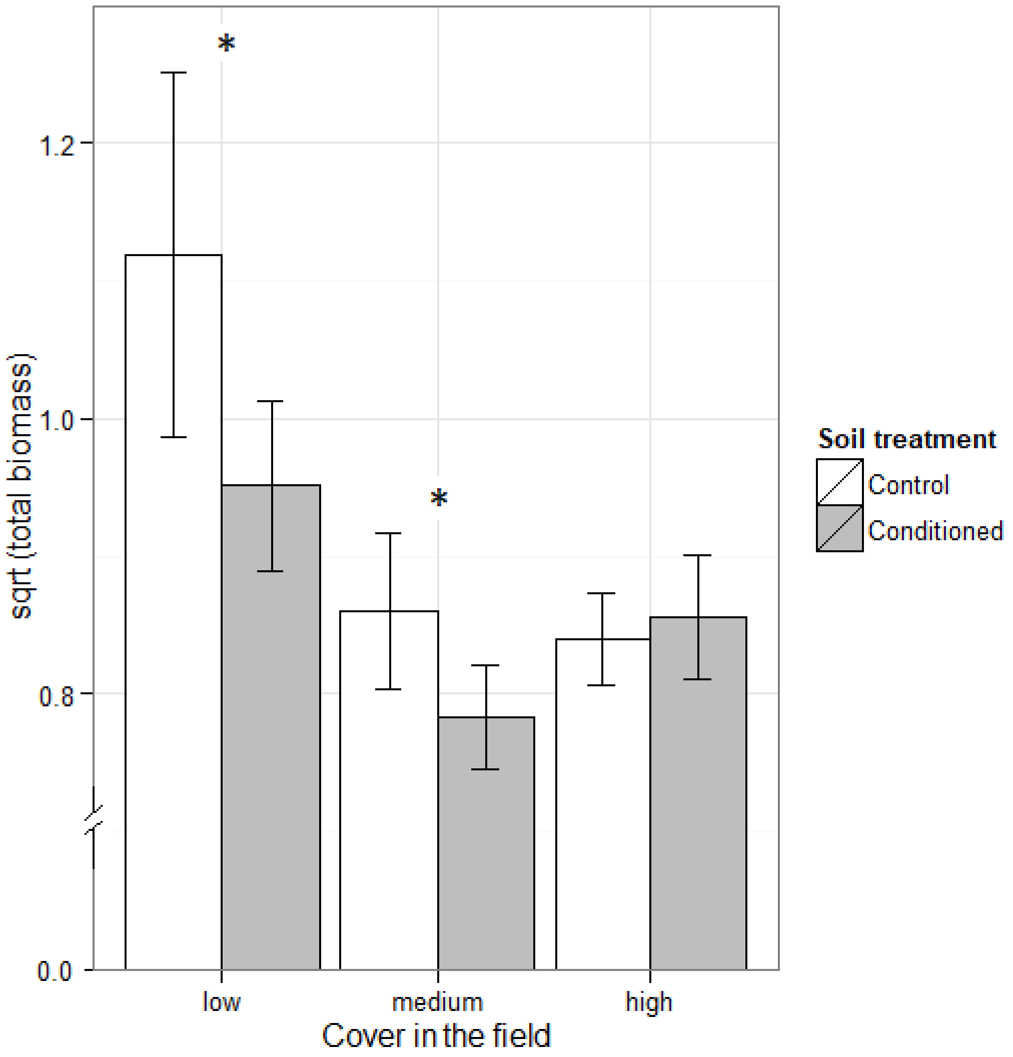


Fig. S3: Square-root transformed biomass of species with low (< 10 %), medium (11-50 %) and high (> 50 %) maximum cover in the field in control and conditioned soil. Asterisks indicate significant ($P < 0.05$) difference between control and conditioned soil. Better performance in conditioned soil compared to control indicates positive PSF, better performance in control soil compared to conditioned soil indicates negative PSF.
